# AlphaFold2: A role for disordered protein prediction?

**DOI:** 10.1101/2021.09.27.461910

**Authors:** Carter J. Wilson, Wing-Yiu Choy, Mikko Karttunen

**Affiliations:** Department of Mathematics, The University of Western Ontario, 1151 Richmond Street, Canada, N6A 5B7; Centre for Advanced Materials and Biomaterials Research, The University of Western Ontario, 1151 Richmond Street, London, Ontario, Canada, N6A 5B7; Department of Biochemistry, The University of Western Ontario, 1151 Richmond Street, Canada, N6A 5C1; Department of Chemistry, The University of Western Ontario, 1151 Richmond Street, Canada, N6A 3K7; Department of Physics and Astronomy, The University of Western Ontario, 1151 Richmond Street, Canada, N6A 5B7

## Abstract

The development of AlphaFold2 was a paradigm-shift in the structural biology community; herein we assess the ability of AlphaFold2 to predict disordered regions against traditional sequence-based disorder predictors. We find that a näaive use of Dictionary of Secondary Structure of Proteins (DSSP) to separate ordered from disordered regions leads to a dramatic overestimation in disorder content, and that the predicted Local Distance Difference Test (pLDDT) provides a much more rigorous metric. In addition, we show that even when used for disorder prediction, conventional predictors can outperform the pLDDT in disorder identification, and note an interesting relationship between the pLDDT and secondary structure, that may explain our observations, and hints at a broader application of the pLDDT to IDP dynamics.

## Introduction

Predicting the three dimensional structure of a protein from its primary amino acid sequence is a grand challenge in molecular structural biology dating back to the late 1950’s ^1,2^. About a year ago in late autumn 2020, AlphaFold2, a deep-learning program, provided a a paradigm-shift in this problem^3^. Not only did it outperform all other groups at the 14th Critical Assessment of protein Structure Prediction (CASP14) ^3^, but it did so with an astonishing accuracy and a large margin, and consequently caused immediate enthusiasm in related fields such as drug development^4^.

The full problem of protein folding is however, multi-faceted, and despite AlphaFold’s stellar success, many problems and open questions remain. As has already been pointed out by several authors ^5–7^, dynamics of protein folding remains a formidable problem; prediction of the folding pathways, effects of mutations, the solution environment, aggregation and, as a very particular category, intrinsically disordered proteins (IDPs).

IDPs remain a major challenge since they are almost entirely devoid of native structure and also because they function primarily as a conformational ensemble ^8–11^ with folding free energy landscapes that are relatively flat^12–14^. This is a direct consequence of their amino acid sequences ^15–17^, in particular the enrichment of disorder-promoting residues over and above order-promoting ones ^18–21^. The application of AlphaFold2 to the prediction of disordered regions and proteins has only briefly been discussed in the literature^6,7,22^, and its performance against traditional predictor methods is currently absent.

In light of the recent publication of the Critical Assessment of protein Intrinsic Disorder (CAID) benchmark ^23^, detailing the performance of over three dozen sequence based disorder predictors and their datasets, we saw an excellent opportunity to benchmark AlphaFold2. Herein we compare the performance of AlphaFold2 to the top performing sequence predictors as determined at CAID. We find that a näive application of structure assignment provided by DSSP ^24^, the primary method for assigning secondary structure based on protein, geometry for the determination of disordered regions, is inaccurate.

The predicted Local Distance Difference Test (pLDDT), which is correlated to the confidence of the structure prediction, provides a better metric for identifying ordered and disordered regions. Furthermore, we find that traditional predictors are capable of outperforming AlphaFold2 in disorder prediction even when the pLDDT is used. We also show how secondary structure and pLDDT scores are interestingly related, providing a potential explanation for the observed performance discrepancy and suggesting a possible link between IDP dynamics and the pLDDT.

## Methodology

### Dataset generation

Two datasets were used in this work, DisProt and DisProt-PDB derived from the DisProt database^25^. Both reference sets are based on the CAID benchmark dataset and are composed of 475 targets, annotated between June 2018 and November 2018 (DisProt release 2018 11). Note that this is less than the 646 targets used at CAID because AlphaFold2 predicted structures do not exist for some sequences. In the DisProt reference set, all residues not labeled as disordered (1) are labelled as ordered (0). In the DisProt-PDB set, residues for which structural data are available are labelled ordered, however a disorder assignment in the DisProt database overrides this order assignment. All residues not covered by either DisProt annotation or PDB structures are masked and were excluded from analysis. As a result the DisProt-PDB dataset contains no ‘uncertain’ residues, all residues considered in this set have either a DisProt annotation or belong to a PDB structure. Additional details pertaining to dataset construction are provided in Supplementary Information and the full list of proteins, structures and combined disorder data are available at https://github.com/SoftSimu/AlphaFoldDisorderData.

AlphaFold2 structures were downloaded from the EMBL database (https://alphafold.ebi.ac.uk/) and run using DSSP^24^ to assign secondary structure. We assume residues belonging to helices, strands, or H-bond stabilized turns are ordered (0) and all other residues are disordered (1). We refer to this as the DSSP predictor or DSSPp for short. We also collected the predicted Local Distance Difference Test (pLDDT) for each structure. Every residue in an AlphaFold2 structure is assigned a value, scaled between 0 and 100, that estimates how well the experimental and predicted structure would agree based on the Local Distance Difference Test (lDDT)^3,22,26^. We transform this value according to the equation,

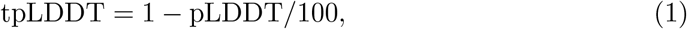

as suggested by Tunyasuvunakool *et al*. ^22^, giving us a pLDDT-based predictor of disorder, where 1 is disordered and 0 is ordered. We refer to this prediction method as the transformed pLDDT or tpLD for short.

We can discretize this pLDDT predictor by classifying a residue with a pLDDT score ≥ *n* as ordered (0) and disordered (1) otherwise; we use pLDDT_*n*_ (or pLD_*n*_ for short), to indicate this binary predictor. Thresholds for *n* were chosen based on the Matthews correlation coefficent (MCC), that has been documented to be an excellent metric for assessing the accuracy of binary classifiers ^27^ and was the approach used at CAID^23^.

The CAID dataset contains predictions made by three dozen predictors; we selected the top 10 performing on the DisProt and DisProt-PDB giving a combined non-redundant set of 11 (fIDPnn^28^, SPOT-Disorder2^29^, RawMSA ^30^, fIDPln ^28^, Predisorder ^31^, AUCpreD ^32^, SPOT-Disorder1^33^, SPOT-Disorder-Single (SPOT-Disorder-S)^34^ , DisoMine ^35^, AUCpreD-np ^32^ and ESpritz-D ^36^). The sequence predictors provide a score between 0 and 1 inclusive as well as a binary disorder/order assignment. No modification to the classification thresholds for these predictors was attempted. Descriptions of disorder prediction methods are provided in the Supplementary Information of the original CAID paper ^23^.

For two vectors *v* and *w* we compute the RMSD as

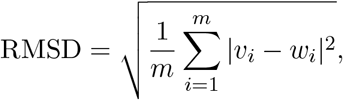

where *m* is the number of elements (residues) in each vector (protein), *v* and *w*. Given binary vectors a random predictor has an RMSD of ∼0.7 on a uniform dataset. Receiver operating characteristic (ROC), area under the curve (AUC), precision-recall, F_1_-score and correlation analysis were all performed using scikit-learn ^37^ and kernel density estimates (KDE) analysis was performed in seaborn ^38^. Descriptions of statistical methods are provided in Supplementary Information.

## Results

### pLDDT performs better than näaive use of DSSP for disorder prediction

Improved performance with tpLD (Eq. 1) over and against DSSPp is evidenced by the ROC curves and AUC values (Figs. 1a, S1a), as well as the precision-recall (PR) curves and F_max_ values (Figs. 1b, S1b) on both the DisProt-PDB and DisProt datasets (Tables S1 and S2). Thresholds for the binary pLD_*n*_ predictor were selected based on the Matthews correlation coefficients which gave values of 76 and 68 for the DisProt and DisProt-PDB datasets respectively (Tables S3 and S4). We refer to these discrete predictors as pLD_76_ and pLD_68_. Unsurprisingly, these values agree with the minimum distance from the ROC curve to the top left of the plot (i.e. (0,1)) (Fig. 1). The difference between these two values undoubtedly stems from the nature of the underlying datasets, while DisProt-PDB contains no uncertain residues, Disprot does. For analysis purposes, we opt to use a combined pLDDT metric, denoted pLD_72_ that is the mean of these two. Data using multiple pLDDT values is provided in Tables S1 and S2.

**Figure 1:**
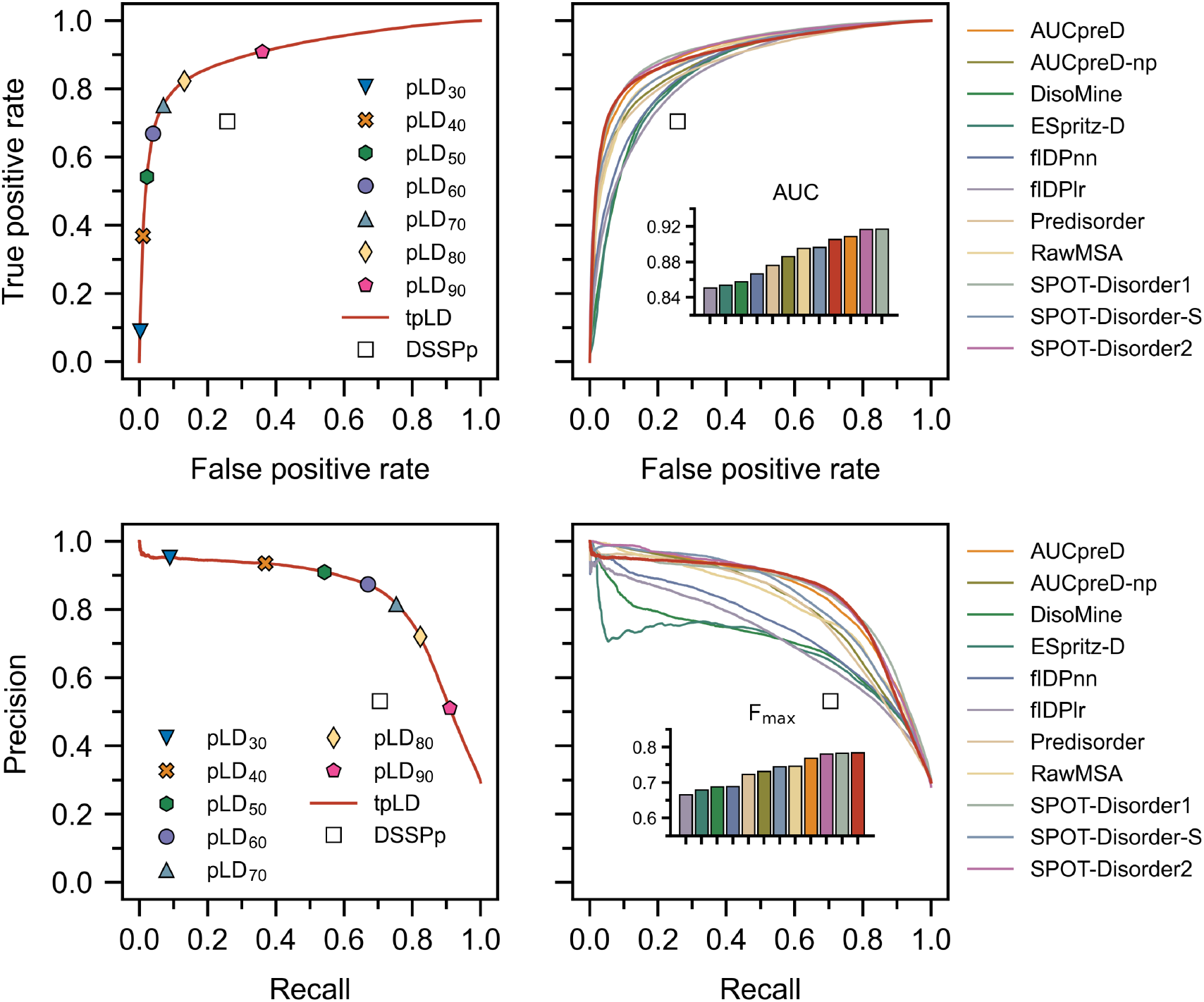
Receiver operating characteristic (ROC) curves (top) and precision-recall (bottom) are depicted for various predictors calculated per-residue on the DisProt-PDB dataset. tpLD (Eq. 1) and various discrete pLD_*n*_ predictors are indicated alongside DSSPp. Inset bar plots show the F_max_ (top inset) and AUC (bottom inset) for the various predictors on the DisProt-PDB dataset (colors correspond to the legend; red is tpLD). The tpLD predictor resulted in one of the highest AUC values and the highest F_max_ on the DisProt-PDB dataset. pLDDT is abbreviated pLD for plotting purposes.

RMSD calculations comparing DSSPp and pLD_72_ demonstrate improved performance for all protein classes, including highly disordered (i.e. *>* 95%) and highly ordered (i.e. *<* 10%), irrespective of dataset (Fig. 2). We note that overall RMSD values are markedly lower for the DisProt-PDB dataset, again likely a result of it lacking “uncertain” residues – residues for which no PDB or experimental data exists. Shifts towards lower RMSD irrespective of dataset, or protein length and disorder content, are also evident for pLD_72_ (Figs. S2, S3). Regression analysis revealed stronger correlations between pLD_72_ and the traditional disorder predictors with respect to residue-wise disorder RMSD when compared with DSSPp (Figs. S4–S7).

**Figure 2:**
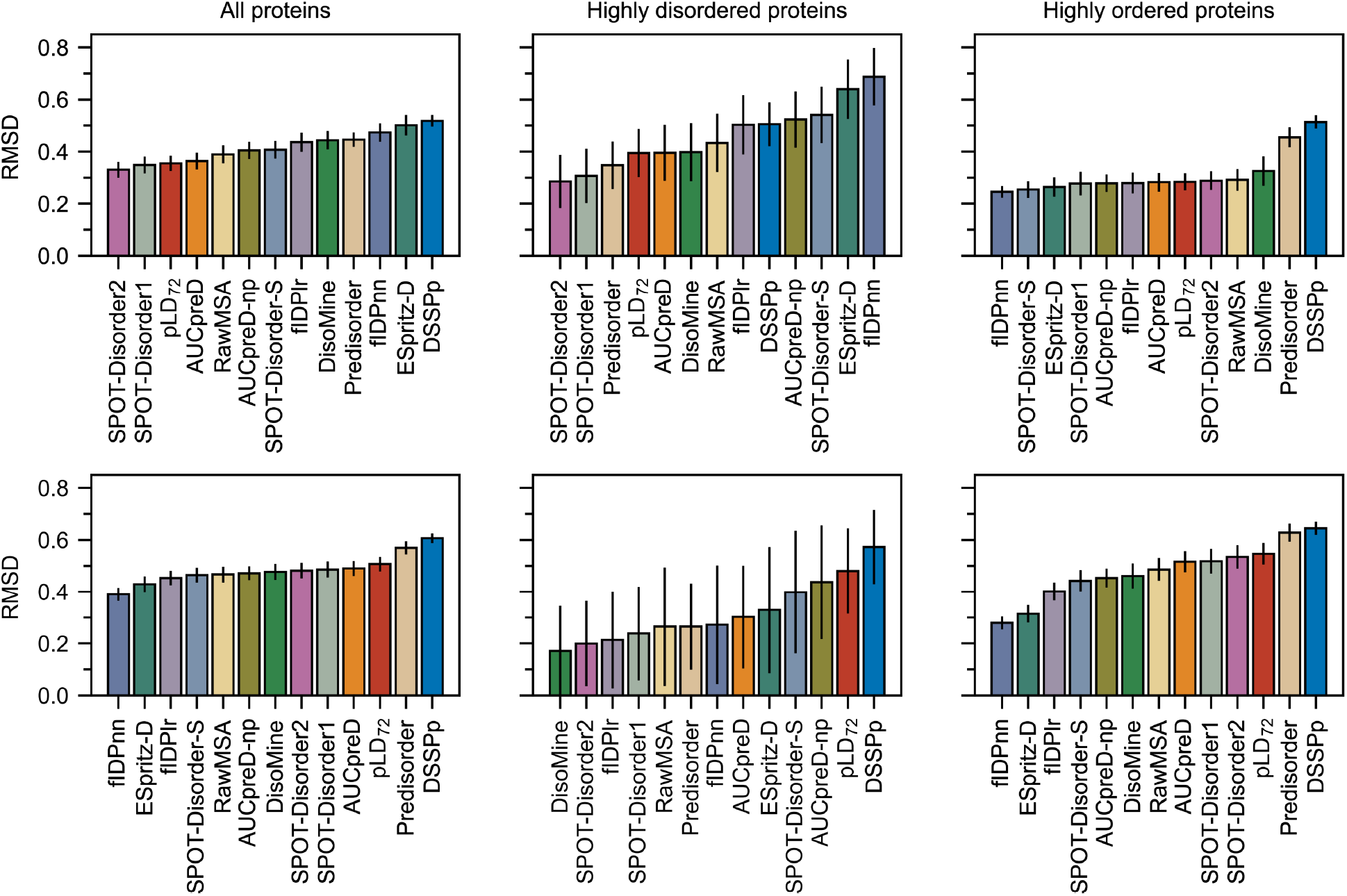
Average RMSD values calculated for the DisProt-PDB (upper) and DisProt (lower) datasets using various prediction methods calculated per-protein. Proteins were assigned to classes (highly disordered i.e. *>* 90% disorder and highly ordered i.e. *<* 10% disorder) based on datasets; specifically with DisProt-PDB, only residues for which PDB or DisProt data were available are considered in the total disorder calculation. Bootstrapping was used to compute averages and estimate errors with 10,000 samples of size 60. pLD_72_ resulted in lower RMSD values on the DisProt-PDB dataset compared to DSSPp, however both showed much higher RMSD on the DisProt dataset. pLDDT is abbreviated pLD for plotting purposes.

Considering global disorder content prediction, we find that on the DisProt dataset, pLD_72_ shows slightly better performance than DSSPp, with a lower mean and a more accurate distribution; however, we note that both methods significantly overestimate disorder content (Fig. 3). On the DisProt-PDB dataset, closer agreement between pLD_72_ and DSSPp is evident based on the mean with both methods returning values similar to experiment. The two distributions are, however, notably different. While that produced by pLD_72_ has a peak around 0.15 in close agreement with experiment, the peak in the distribution produced by DSSPp is larger and shifted to a higher value around 0.3. This is all to say that a näive application of DSSP for the prediction of disordered and ordered regions for AlphaFold2 structures, specifically the assumption that helical and strand regions are ordered and coiled regions are unstructured, leads to poorer prediction (i.e., higher RMSD, lower AUC and higher F_max_) of disordered regions and an overestimation in disorder content.

**Figure 3:**
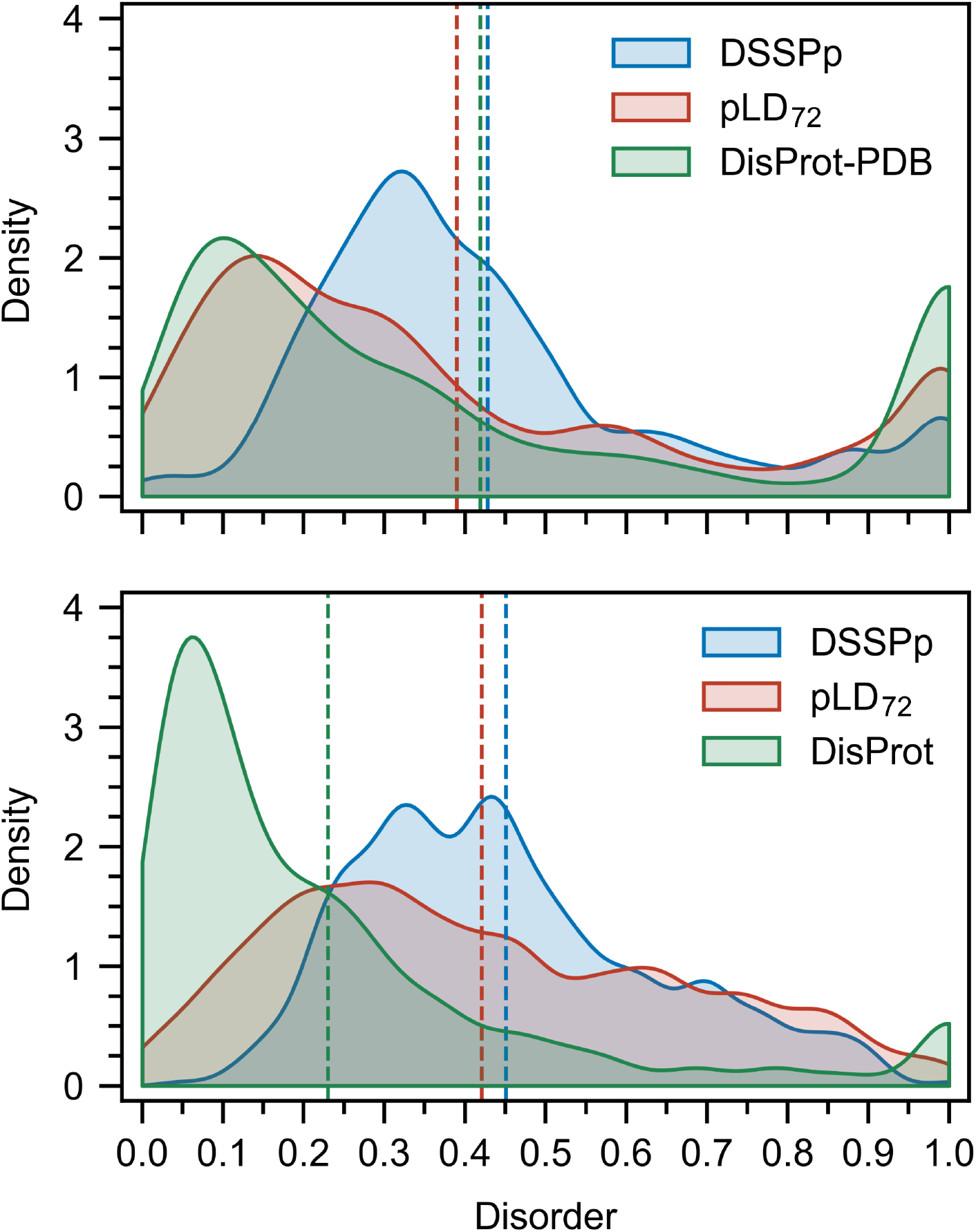
Distribution of disorder content per-protein in the DisProt-PDB and DisProt datasets depicted alongside the distributions predicted by pLD_72_ and DSSPp. Bin-widths were set at 0.5 and bootstrapping was used to compute the distributions and average values (vertical dashed lines) with 10,000 samples of size 60. On the DisProt-PDB dataset close agreement between experiment and pLD_72_ is evident; conversely, on both the DisProt-PDB and DisProt datsets, DSSPp predicted a higher disorder content. pLDDT is abbreviated pLD for plotting purposes.

### Sequence predictors can still outperform AlphaFold2 on disorder prediction

Comparing the pLDDT-based and DSSPp predictors to various sequence-based predictors revealed performance differences amongst the methods. Notably, tpLD (Eq. 1) performed exceptionally well on the DisProt-PDB dataset posting the largest F_max_ (0.784) and one of the largest AUC (0.905) values of the methods considered (Fig. 1, Tables S1 and S3). This was also evidenced by pLD_72_ which had the highest MCC (0.701) (Table. S1) and one of the lowest RMSD values (Fig. 2) on the Disprot-PDB dataset. Interestingly, on the DisProt dataset, both tpLD (Eq. 1) and DSSPp performed significantly worse and were readily outperformed by the other predictor methods, in particular fIDPnn (F_max_: 0.357 (DSSPp), 0.429 (tpLD), 0.457 (fIDPnn); AUC: 0.635 (DSSPp), 0.731 (tpLD), 0.794 (fIDPnn)), which outperformed all other predictors, as evidenced by the ROC, PR, and RMSD analyses. We note that with respect to MCC, pLD_72_ still performed well on both the DisProt and DisProt-PDB datasets achieving scores of 0.310 and 0.697 respectively (Tables S1, S2). In agreement with the CAID results we found that SPOT-Disorder2, fIDPnn, RawMSA and AUCpreD all performed exceptionally well (Figs. 1 and S1, Tables S3 and S4) ^23^.

### Secondary structure codons (SSC) reveal relationships between the pLDDT and secondary structure

In order to explain the discrepancy between the pLDDT-based and DSSP predictors with respect to local and global disorder prediction, we considered how pLDDT values were assigned to the secondary structures. Kernel density estimates (KDE) of the distribution of pLDDT values sampled over all residues reveal a strong left-skew for all but the coil secondary structure which exhibits a right-skewed bimodal distribution with peaks around 94 and 35 (Fig. 4). Residues assigned to *β*-strand and *β*-bridge structures are the most likely to be assigned to large pLDDT values, followed by helical and H-bond stabilized turns. To provide a more detailed picture of the distributions, we introduce the concept of a secondary structure codon (SSC), a triplet describing the local secondary structure at a given residue. Analysis of the distributions of pLDDT values for each SSC revealed that residues predicted to belong to both the ends (HHC/CHH/HHT/THH) and middle (HHH) of helices can have pLDDT values *<*50 (Fig. S8), this was not observed for residues belonging to the middle (EEE) and ends of *β*-strands(EEC/CEE/EET/TEE) (Fig. S9). For highly coiled residues (CCC/CCT/TCC), both high (*>* 80) and low (*<* 50) pLDDT values were observed (Figs. S10 and S11).

**Figure 4:**
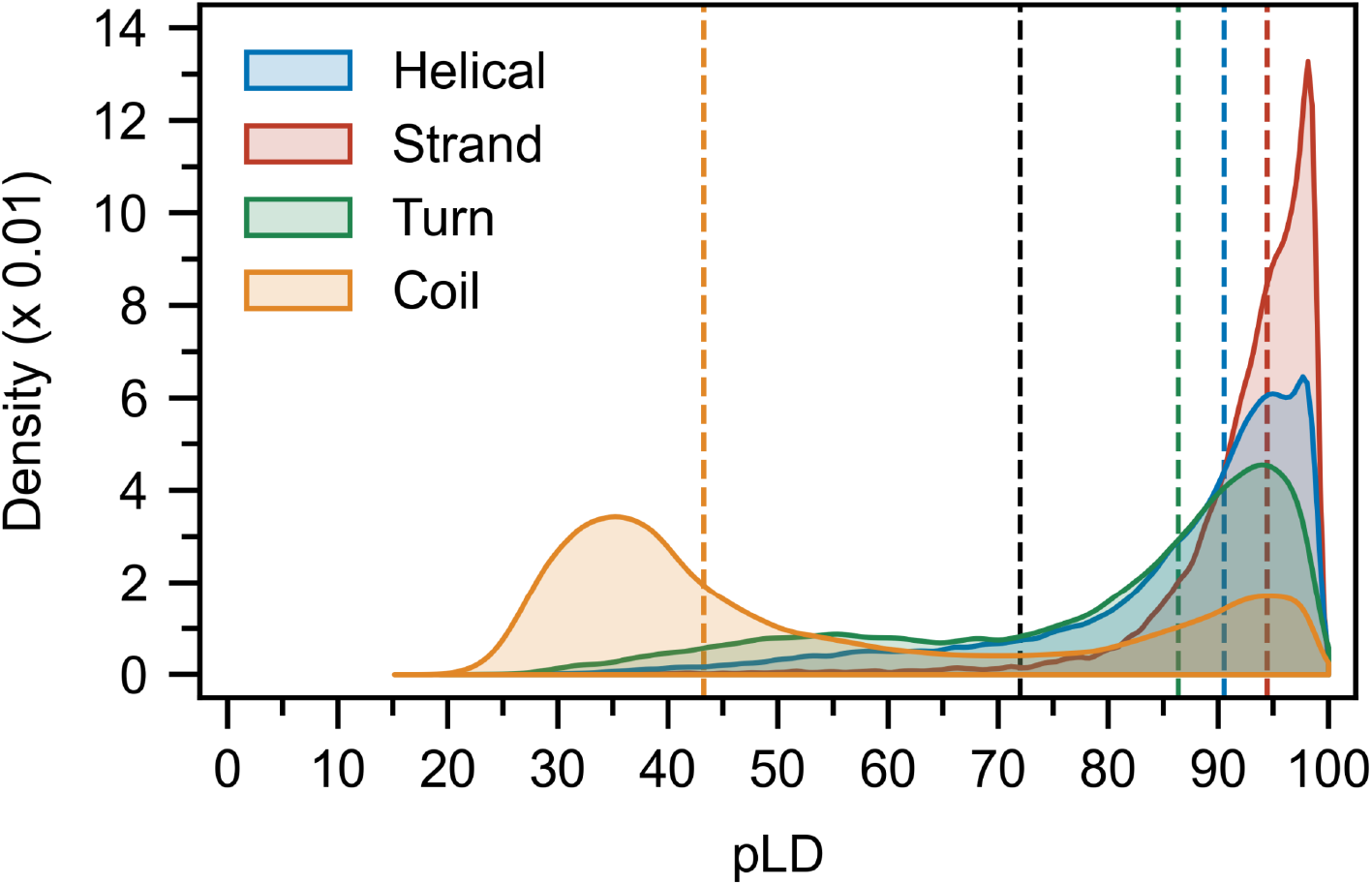
Distribution of pLDDT values per-residue calculated for each secondary structure class. Bin-widths were set at 0.5 and bootstrapping was used to compute the distributions and mean values (colored vertical dashed lines; black dashed line represents pLD_72_) with 10,000 samples of size 500. A bimodal distribution is evident for the coil structures, and while strand, helical and turn regions are on average assigned to high pLDDT values, residues belonging to each can sample much lower values. pLDDT is abbreviated pLD for plotting purposes.

## Discussion

AlphaFold2 has been a paradigm-shift in structural biology, providing a tentative solution to the protein folding problem that has persisted over half a century^1^. Since the time that problem was posed by Perutz and Kendrew, a new class of proteins has been discovered and IDPs have become the focus of much study ^8,9,39,40^. Over the past two decades much effort has been devoted to developing methods for identifying disordered regions given the primary sequence of a protein ^23,41–45^, herein, we assess the applicability of AlphaFold2 to this problem.

We find, and strongly stress, that simply inferring a residue in an AlphaFold2 structure assigned by DSSP to a helical, strand, or H-bond stabilized turn is ordered, and otherwise is disordered, results in an overestimation of disorder content and a poor prediction of disordered regions. Instead, employing the pLDDT, a measure of the expected position error at a given residue and originally purposed to assess interdomain accuracy, provides a much more accurate metric for determining global and local disorder content. Using the pLDDT as a disorder predictor metric we observe impressive performance on the DisProt-PDB dataset when compared to conventional disorder predictors (Fig. 1). While poorer performance is observed on the plain DisProt dataset, pLDDT does outperform naïve use of DSSP in both cases.

Secondary structure and global disorder analyses point to a potential root of the prediction discrepancy between pLDDT and DSSP, simply put, for AlphaFold2, not all secondary structures are created equal. AlphaFold2 will readily assign a coiled geometry and a high pLDDT value to the same residue, and conversely assign low pLDDT values to structured regions (Fig. 4). While a DSSP predictor assumes that coils are disordered and helices are ordered, a pLDDT predictor will account for the fact that a coil may be more ordered and a helix more disordered for certain residues in certain proteins. It is this former case that is likely resulting in the improved performance observed for pLDDT and underscores the importance of the nuance provided by this metric for disordered protein prediction.

Second to the problem of predicting the (dis)orderedness of a region within a protein is predicting the structural dynamics and transitions (i.e. order-to-disorder, disorder-to-order, disorder-to-disorder) that an IDP may undergo^41,46^. In light of the secondary structure analysis, the pLDDT may be just such a means for extracting this information, namely the transientness of secondary structures, their potential for transition upon binding and their functional importance. A helix with a low pLDDT may be more transient, existing frequently in a disordered, unfolded state, than a helix with a high pLDDT and conversely, a coiled region with a high pLDDT, may suggest a disorder-order transition and/or its conserved role in some biophysical interaction. We here reiterate that by their very nature IDPs exhibit a high degree of conformational flexibility, allowing them to interact with multiple binding partners in a variety of ways ^47–53^. While it is the case that a single, static, AlphaFold2 structure cannot adequately describe these often large conformational ensembles ^8–10^, the ability of the program to predict with relatively high accuracy the location of disordered regions is nonetheless impressive, and refinement of the training set to account for more accurate disordered structures could further improve performance. In addition, thorough analysis of the pLDDT score as it relates to disorder-order transitions, as well as the local function and dynamics of IDP motifs, may further enhance the utility of AlphaFold2 to the IDP community.

While experimental NMR ^54–63^, and high-quality molecular simulations ^64–75^ are some of the most accurate methods for determining the (dis)ordered nature and dynamics of proteins, fast and computationally efficient methods play an important role. Unlike conventional predictors however, AlphaFold2 supplies both a pLDDT score, that can provide an accurate prediction of protein disorderedness, in addition to a three-dimensional structure, that when taken in tandem, may also provide insight into the underlying dynamics of disordered protein regions.

## Conclusion

In this study, we have assessed the ability of AlphaFold2 to predict disordered protein regions. We benchmark the program on the DisProt-PDB and DisProt datasets developed for CAID, and find it to perform quite well, exceeding the performance of 11 traditional predictors on the DisProt-PDB dataset. Furthermore, we observe that the pLDDT score assigned to each residue by AlphaFold2 provides an impressive metric for assessing disorder, far surpassing a näaive application of DSSP. Our analysis also reveals a link between secondary structure and the pLDDT score, suggesting that continued research into this metric may reveal a fundamental connection to the dynamics of disordered proteins.

## Acknowledgements

The authors thank SharcNet and Compute Canada for computational resources.

