## Supplementary Information for "AlphaFold2: A role for disordered protein prediction?"

### Supplemental Methods

#### Dataset generation

$$P = \begin{bmatrix} A & M & K & L & A & T & E & T & G & E & A & H & L & K & A & M \\ - & - & 1 & 1 & 1 & 1 & 1 & - & - & - & - & - & - & - & - & - \\ - & - & - & - & - & - & S & S & S & S & S & S & S & - & - & - \\ 0 & 0 & 1 & 1 & 1 & 1 & 1 & 0 & 0 & 0 & 0 & 0 & 0 & 0 & 0 & 0 \\ - & - & 1 & 1 & 1 & 1 & 1 & 0 & 0 & 0 & 0 & 0 & 0 & - & - & - \end{bmatrix}$$

$P$ represents a single protein and some of the data used in the analysis. The first row is the amino acid sequence, the second row is the DisProt annotation (i.e. 1 = disordered, 0 = ordered, - = no data), the third row is the PDB annotation (i.e. - = no structural data present, S = structural data present), the fourth row is the DisProt dataset entry for this protein, the fifth row is the DisProt-PDB dataset entry for this protein. Note how residues with no DisProt data are assumed to be structured in the DisProt dataset entry while only residues for which either PDB or DisProt or both is available are included in the DisProt-PDB dataset entry. Also note that the residue 6 in the middle structured region is assigned to disorder in the DisProt-PDB, this is because disorder supercedes structure present in the PDB database.

#### Statistical analysis methods

**True positive rate** (TPR): also called the sensitivity, this is the proportion of positive classifications that are actually positive, given by $TPR = \left(\frac{TP}{TP+FN}\right)$.

**False positive rate** (FPR): also called the fall-out, this is the proportion of positive classifications that are actually negative, given by $FPR = 1 - TPR$.

**True negative rate** (TNR): also called the specificity, this is the proportion of negative

Table 1: Confusion matrix (also known as error matrix or matching matrix depending on the context) for the disorder prediction problem. A given residue is experimentally classified as disordered or ordered, and the prediction can either classify a residue as disordered or ordered. This gives rise to four classification scenarios: true positive (TP), false positive (FP), false negative (FN) and true negative (TN).

| | | Experiment | |
|---|---|---|---|
| | | Disordered | Ordered |
| Predictor | Disordered | $TP$ | $FP$ |
| | Ordered | $FN$ | $TN$ |

classifications that are actually negative, given by $TNR = \left(\frac{TN}{TN+FP}\right)$.

**False negative rate** (FNR): also called the fall-out, this is the proportion of negative classifications that are actually positive, given by $FNR = 1 - TNR$.

**Reciever operating characteristic**[1] (ROC) and **precision-recall** (PR) are both methods for analyzing the data encoded in the confusion matrix (Table 1) allowing one to construct a point either in ROC or PR space. In ROC space, we plot the $FPR$ vs $TPR$. Conversely, in PR space we plot the Recall ($TPR$) vs Precision $\left(\frac{TP}{TP+FP}\right)$. We note that because the Recall and Precision equations lack true negatives PR curves can can perform poorly if the underlying dataset is imbalanced with few negatives; the ROC curve is insensitive to class imbalance because it does account for these true negatives.

**Precision/Positive predictive value** (PPV): the proportion of positive results that are actually true positives $\left(\frac{TP}{TP+FP}\right)$.

**Area under curve** (AUC): the numerically integrated (trapezoidal) area under the ROC curve that provides an aggregate measure of performance across all possible classification thresholds.

$\mathbf{F}_{\max}$: the maximum F1 score[2], which is itself the the harmonic mean of the precision $\left(\frac{TP}{TP+FP}\right)$ and recall ($TPR$) and takes into account predictions across the entire sensitivity spectrum[3].

**Balanced accuracy** (BAC): the average of the sensitivity $\left(\frac{TP}{TP+FN}\right)$ and specificity $\left(\frac{TN}{FP+TN}\right)$

that is particularly useful when the underlying classes are imbalanced.

**Matthews correlation coefficient**[4] (MCC): the Pearson product-moment correlation coefficient between the actual and predicted values. Given explicitly from the confusion matrix (Table 1) as

$$\mathrm{MCC} = \frac{TP \times TN - FP \times FN}{\sqrt{(TP + FP)\,(TP + FN)\,(TN + FP)\,(TN + FN)}}.$$

Unlike the $F_1$ score and the accuracy the MCC considers all four classes of the confusion matrix making it arguably a superior metric especially when dealing with unbalanced data[5].

### Supplemental Figures and Tables

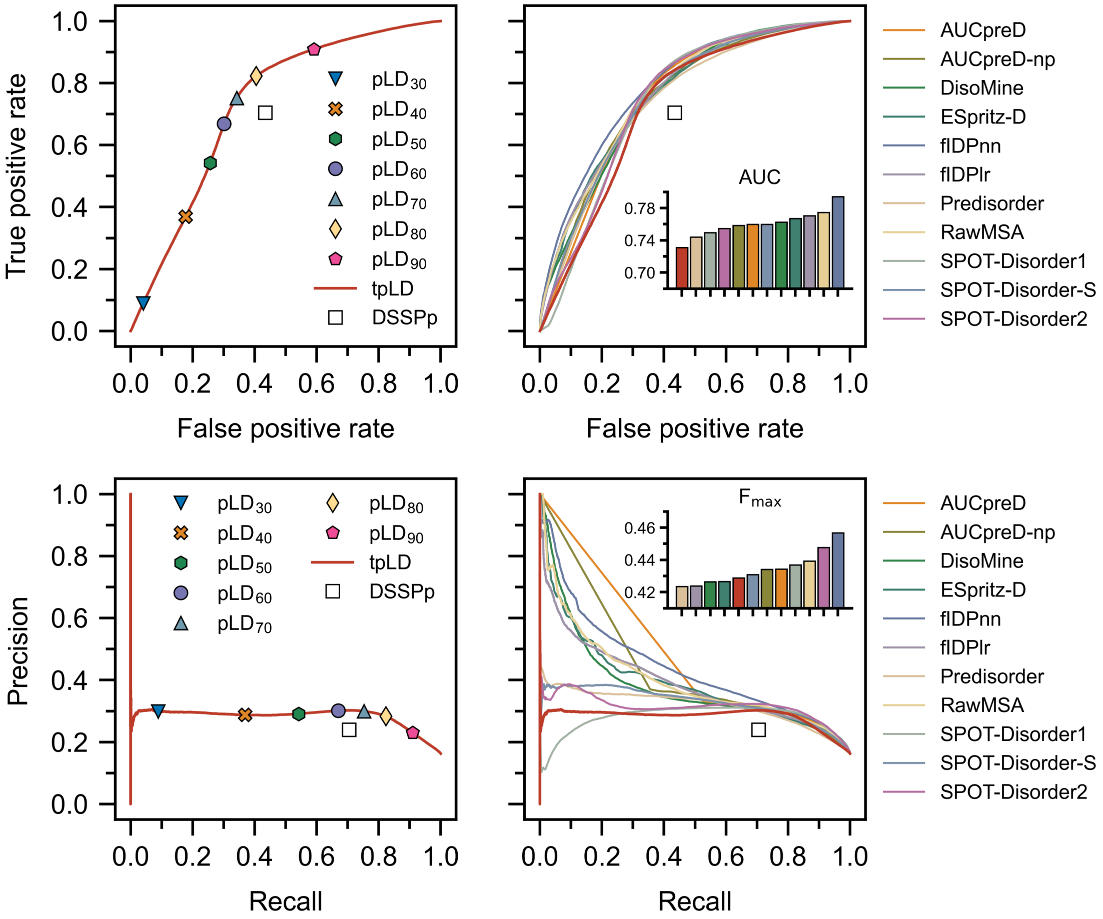


Figure S1: Receiver operating characteristic (ROC) curves (top) and precision-recall (bottom) are depicted for various predictors calculated per-residue on the DisProt dataset. tpLD (Eq. 1 in main text) and various discrete $pLD_n$ predictors are indicated alongside DSSPp. Inset bar plots show the $F_{max}$ (top inset) and AUC (bottom inset) for the various predictors on the DisProt dataset (colors correspond to the legend; red is tpLD). pLDDT is abbreviated pLD for plotting purposes.

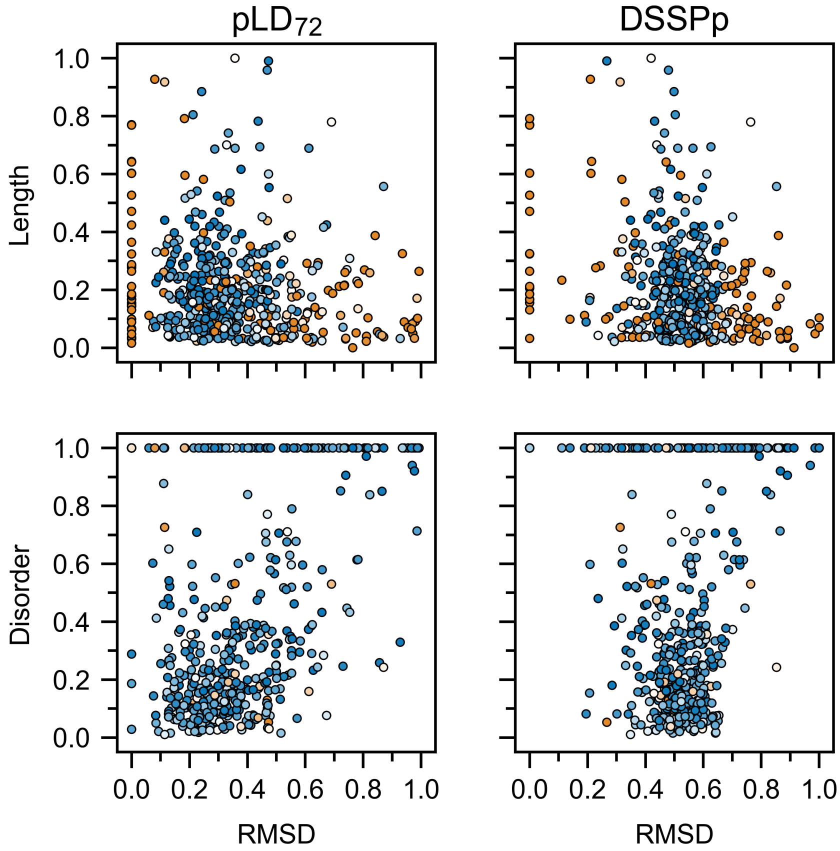


Figure S2: Relationship between protein length (scaled between 0 and 1) and RMSD and protein-wise disorder content and RMSD compared between $pLD_{72}$ and DSSPp on the DisProt-PDB dataset. Colors (0: blue, 0.5: white, 1: orange) indicate disorder content (upper plots) or length (scaled between 0 and 1) (lower plots). Each point represents a protein. pLDDT is abbreviated pLD for plotting purposes.

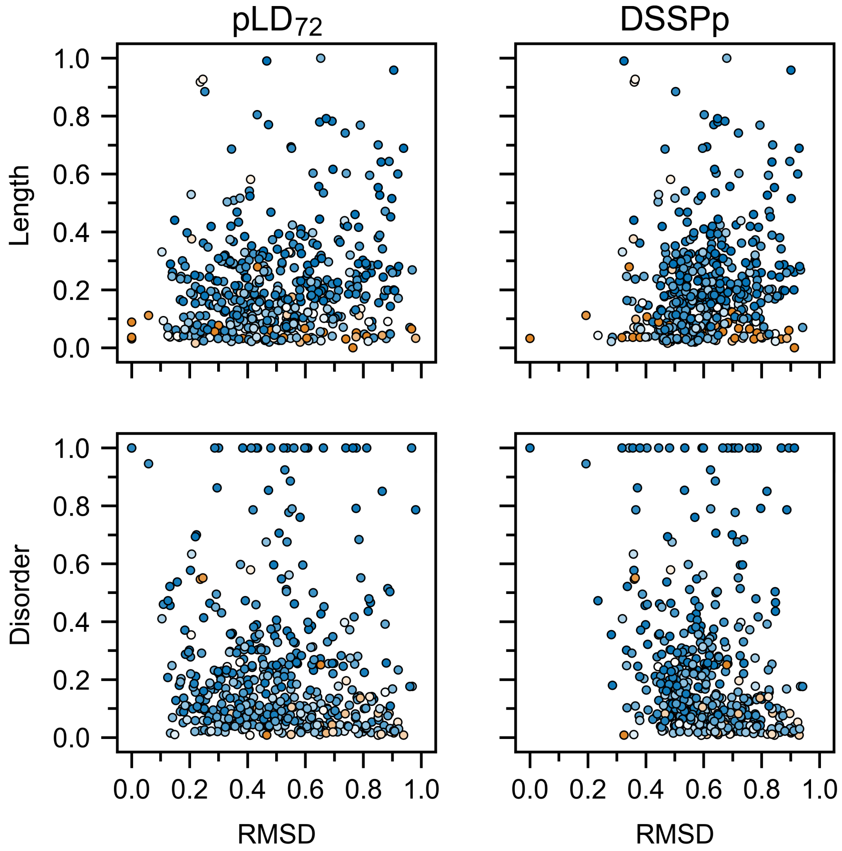


Figure S3: Relationship between protein length (scaled between 0 and 1) and RMSD, and protein-wise disorder content and RMSD compared between pLD$_{72}$ and DSSPp on the DisProt dataset. Colors (0: blue, 0.5: white, 1: orange) indicate disorder content (upper plots) or length (scaled between 0 and 1) (lower plots). Each point represents a protein. pLDDT is abbreviated pLD for plotting purposes.

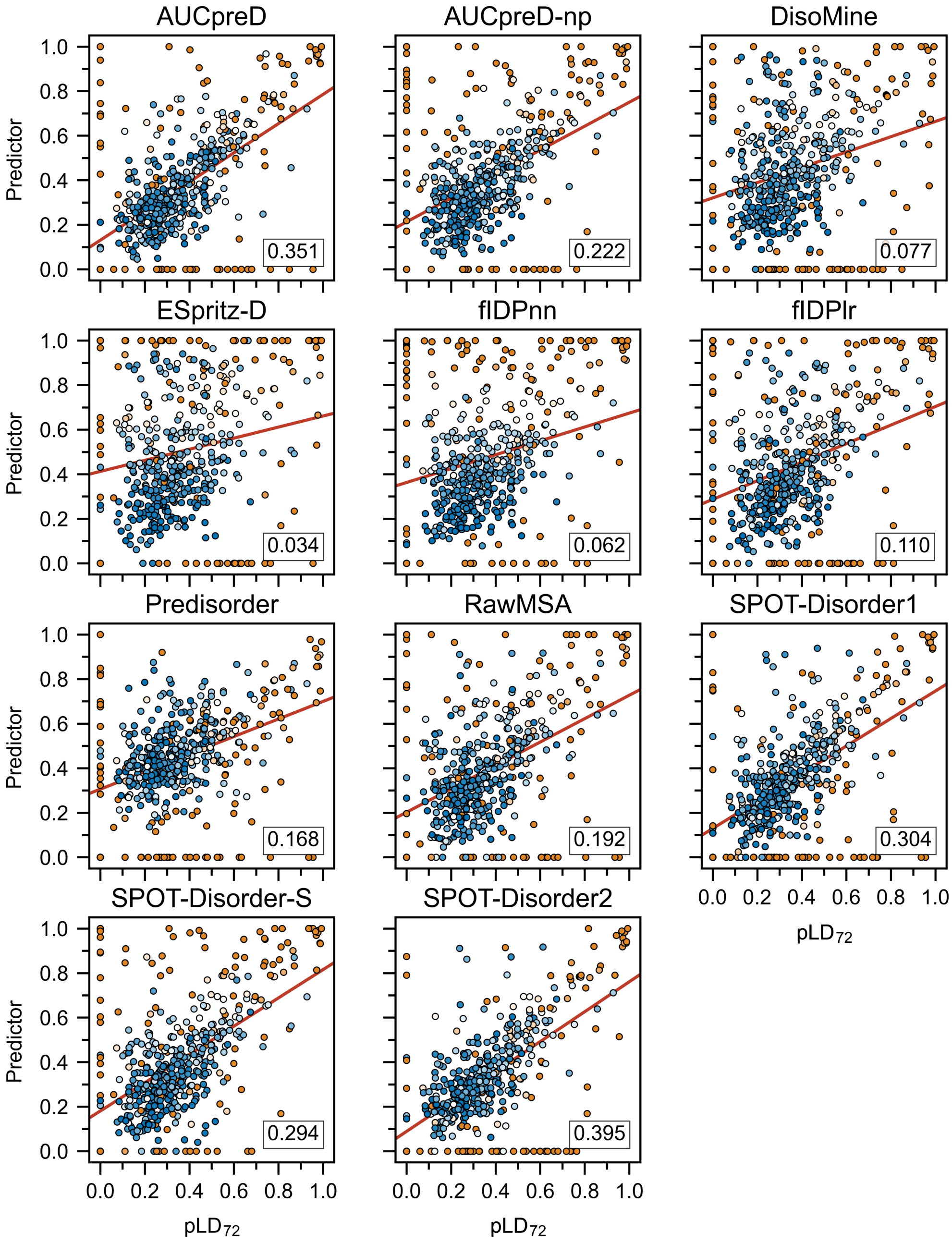


Figure S4: Correlation between conventional predictors and $pLD_{72}$ for predicting protein-wise disorder RMSD on the DisProt-PDB dataset. Colors (0: blue, 0.5: white, 1: orange) indicate protein length (scaled between 0 and 1). Regression lines are colored red and the number in the bottom right is the $r^2$. Each point represents a protein. pLDDT is abbreviated pLD for plotting purposes.

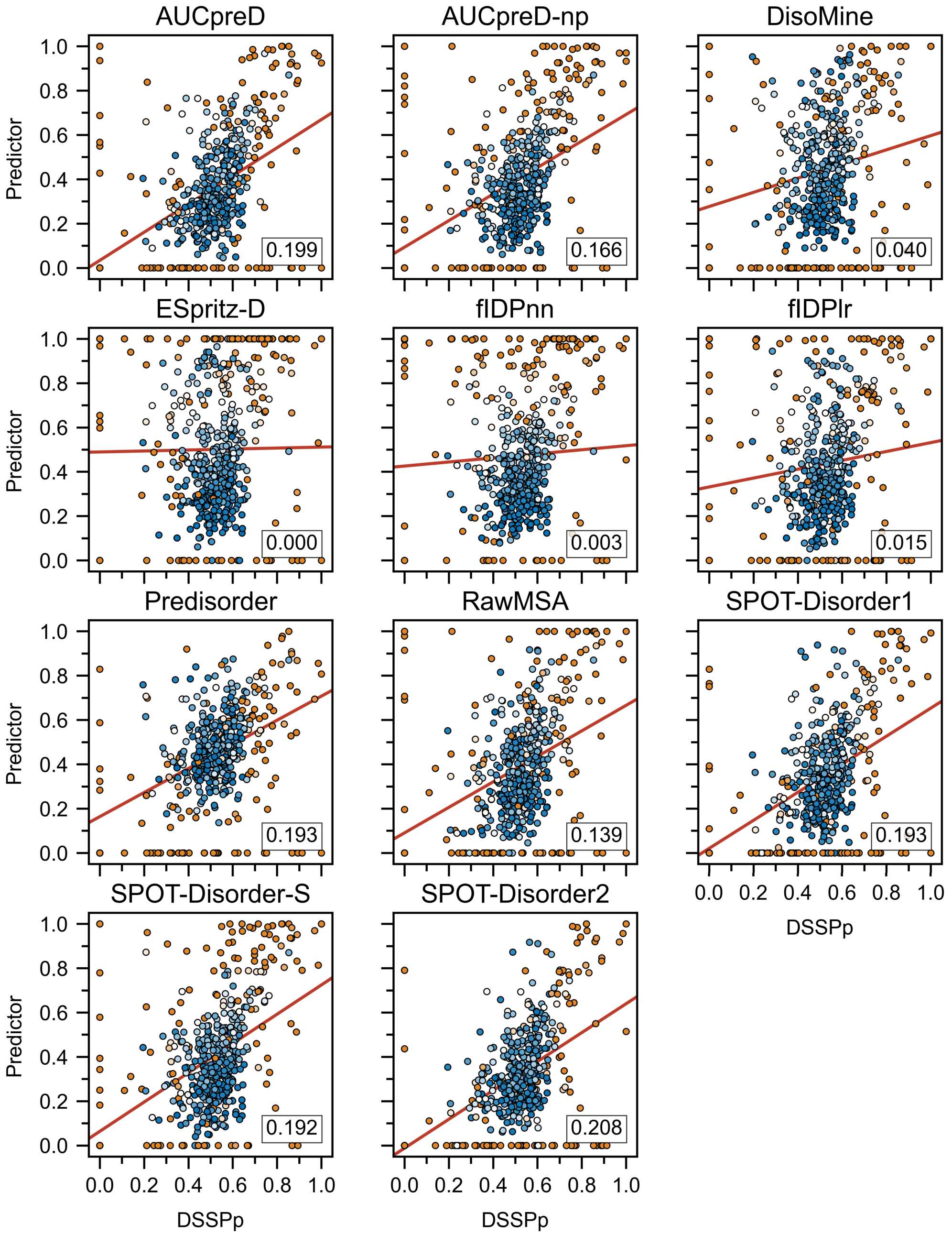


Figure S5: Correlation between conventional predictors and DSSPp for predicting protein-wise disorder RMSD on the DisProt-PDB dataset. Colors (0: blue, 0.5: white, 1: orange) indicate protein length (scaled between 0 and 1). Regression lines are colored red and the number in the bottom right is the $r^2$. Each point represents a protein.

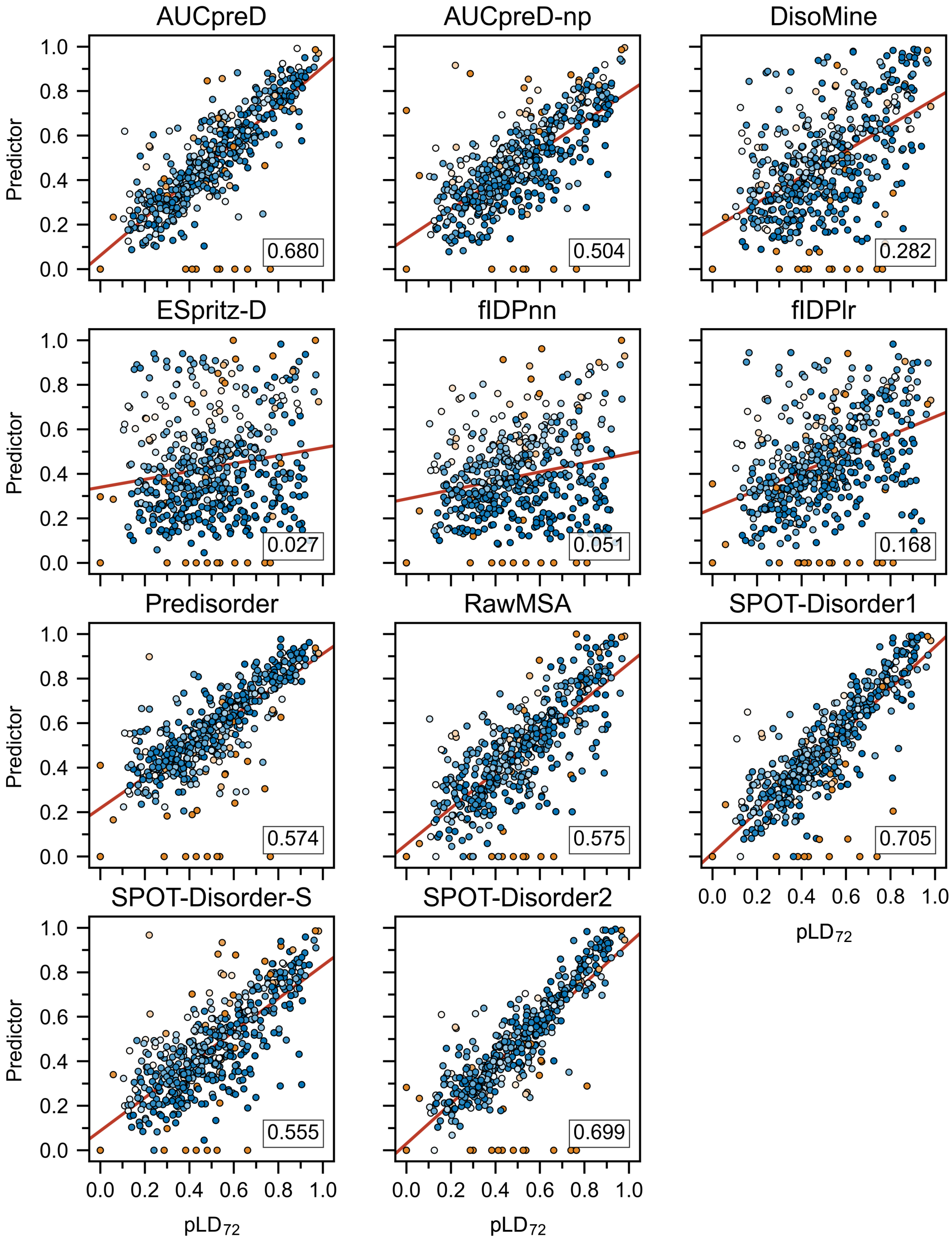


Figure S6: Correlation between conventional predictors and $pLD_{72}$ for predicting protein-wise disorder RMSD on the DisProt dataset. Colors (0: blue, 0.5: white, 1: orange) indicate protein length (scaled between 0 and 1). Regression lines are colored red and the number in the bottom right is the $r^2$. Each point represents a protein. pLDDT is abbreviated pLD for plotting purposes.

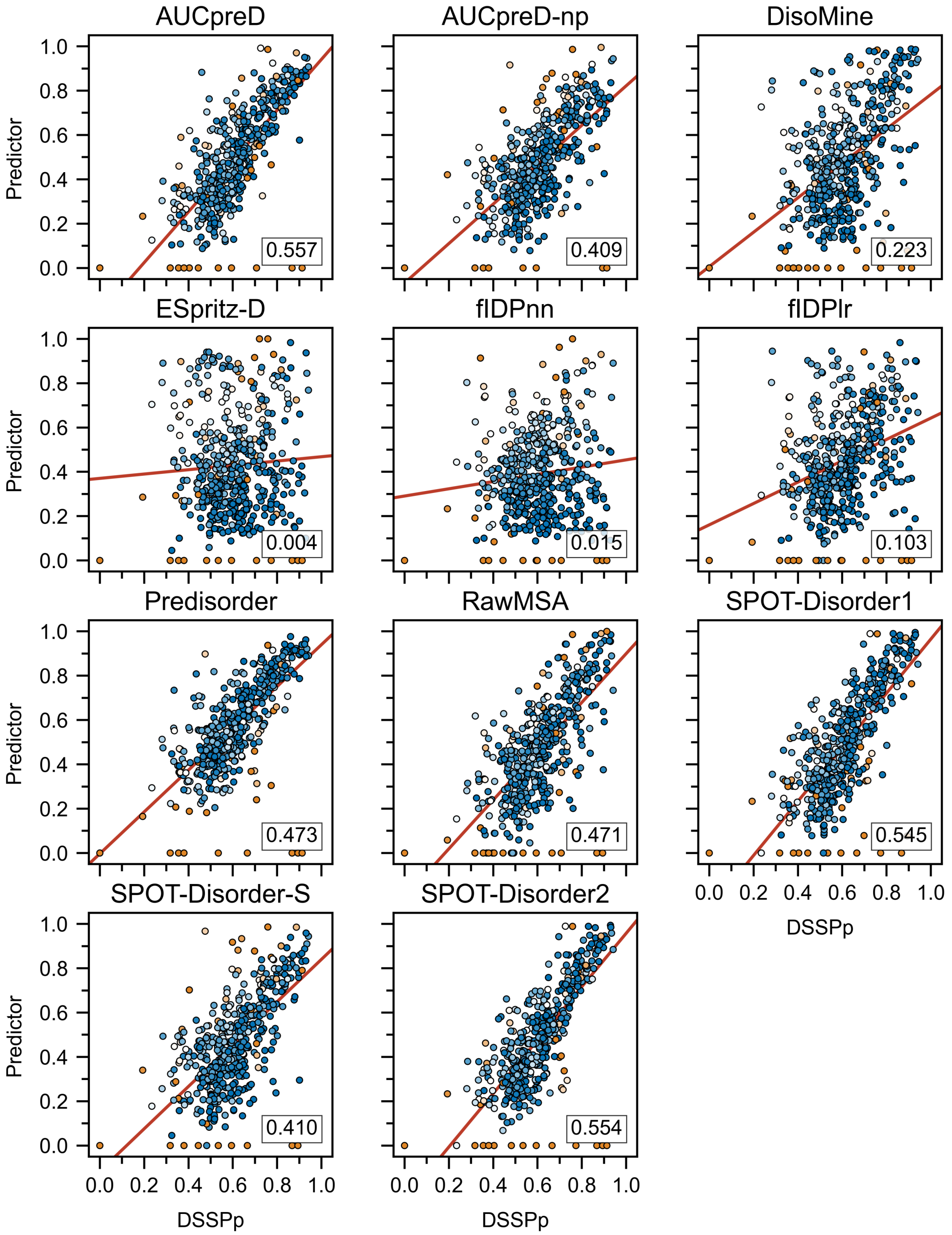


Figure S7: Correlation between conventional predictors and DSSPp for predicting protein-wise disorder RMSD on the DisProt dataset. Colors (0: blue, 0.5: white, 1: orange) indicate protein length (scaled between 0 and 1). Regression lines are colored red and the number in the bottom right is the $r^2$. Each point represents a protein.

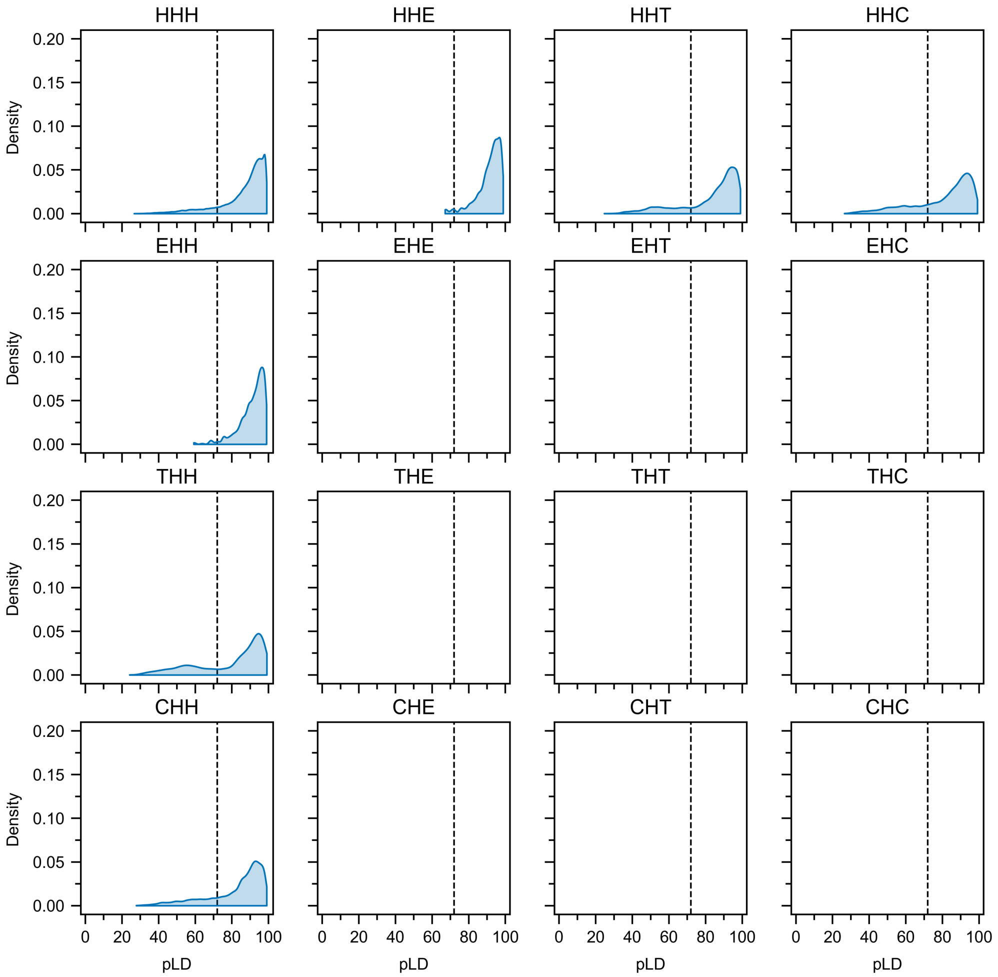


Figure S8: Distributions of pLD values for helical ($\alpha$-helix, 310-helix and $\pi$-helix) secondary structure codons (SSC). Bin-widths were set at 0.5, vertical lines indicate $\mathrm{pLD}_{72}$. pLDDT is abbreviated pLD for plotting purposes.

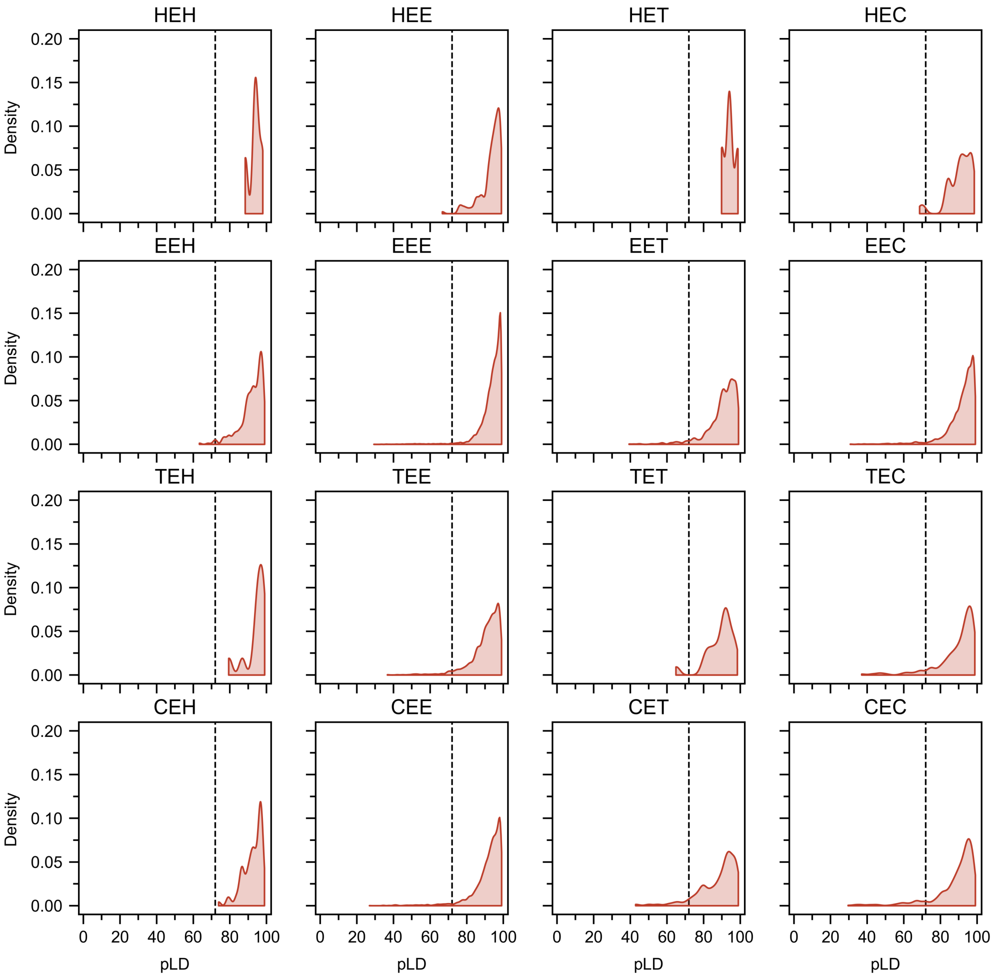


Figure S9: Distributions of pLD values for strand ($\beta$-strand and $\beta$-bridge) secondary structure codons (SSC). Bin-widths were set at 0.5, vertical lines indicate $pLD_{72}$. pLDDT is abbreviated pLD for plotting purposes.

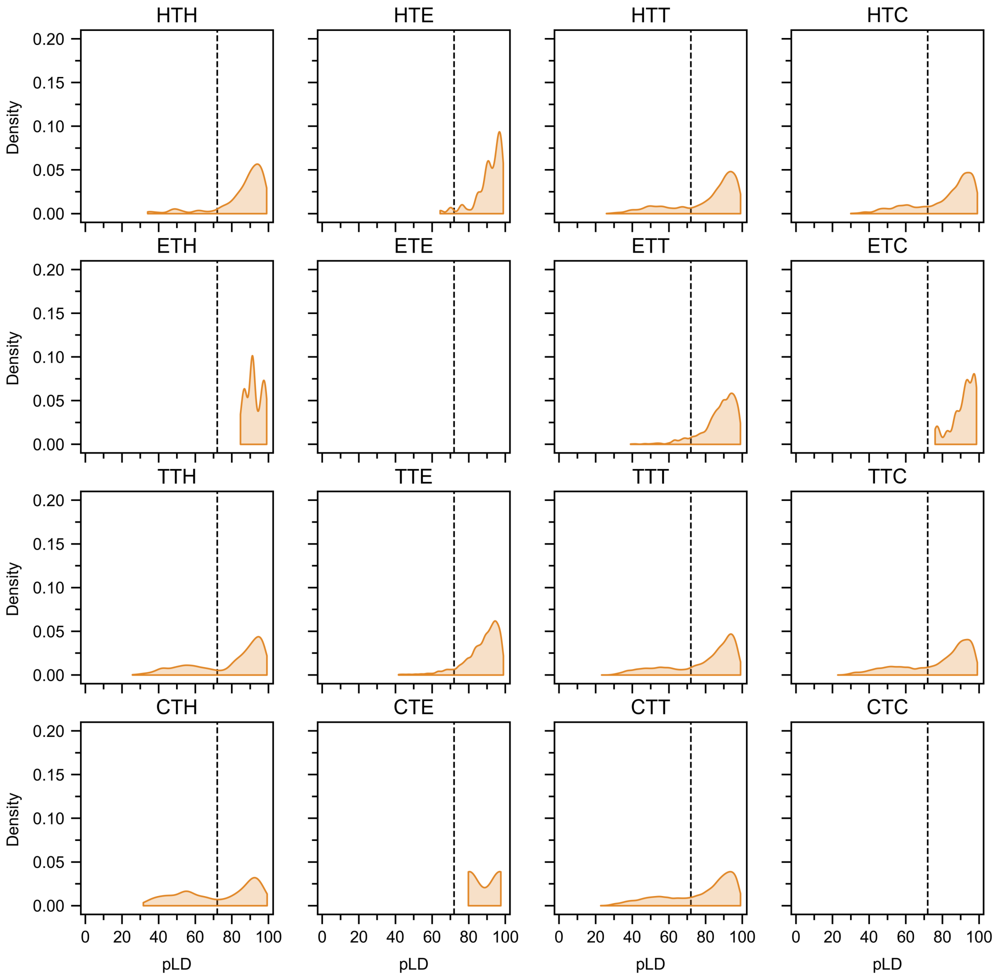


Figure S10: Distributions of pLD values for turn (H-bond stabilized turn) secondary structure codons (SSC). Bin-widths were set at 0.5, vertical lines indicate $pLD_{72}$. pLDDT is abbreviated pLD for plotting purposes.

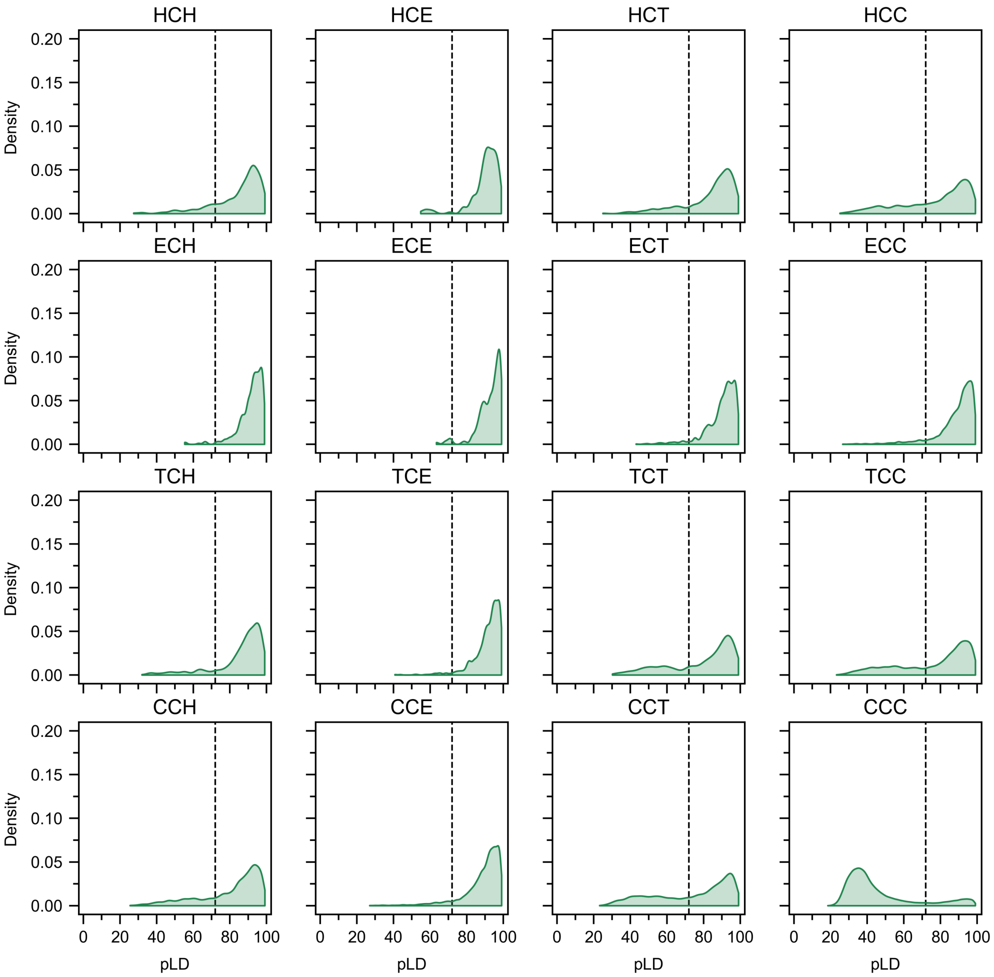


Figure S11: Distributions of pLD values for coil (coil or bend) secondary structure codons (SSC). Bin-widths were set at 0.5, vertical lines indicate $\text{pLD}_{72}$. pLDDT is abbreviated pLD for plotting purposes.

Table S1: Tabulated performance metrics for pLD-based (tpLD and $pLD_n$) and DSSPp predictors on the DisProt-PDB dataset. Abbreviations: Matthews correlation coefficent (MCC); area under ROC curve (AUC); maximum F1-score ($F_{max}$); true postivie rate (TPR); false positive rate (FPR); true negative rate (TNR); positive predictive value (PPV); balanced accuracy (BAC); number of proteins covered (COV); predicted Local Distance Difference Test (pLD).

| Pred | MCC | AUC | $F_{max}$ | TPR | FPR | TNR | PPV | BAC | COV |
|---|---|---|---|---|---|---|---|---|---|
| DSSPp | 0.418 | 0.724 | 0.606 | 0.704 | 0.257 | 0.743 | 0.532 | 0.724 | 478 |
| tpLD | N/A | 0.905 | 0.784 | N/A | N/A | N/A | N/A | N/A | 478 |
| $pLD_{30}$ | 0.243 | 0.544 | 0.453 | 0.089 | 0.002 | 0.998 | 0.952 | 0.544 | 478 |
| $pLD_{32}$ | 0.308 | 0.569 | 0.453 | 0.141 | 0.003 | 0.997 | 0.948 | 0.569 | 478 |
| $pLD_{34}$ | 0.368 | 0.598 | 0.453 | 0.200 | 0.005 | 0.995 | 0.944 | 0.598 | 478 |
| $pLD_{36}$ | 0.422 | 0.626 | 0.453 | 0.259 | 0.007 | 0.993 | 0.942 | 0.626 | 478 |
| $pLD_{38}$ | 0.469 | 0.654 | 0.474 | 0.317 | 0.009 | 0.991 | 0.937 | 0.654 | 478 |
| $pLD_{40}$ | 0.510 | 0.679 | 0.529 | 0.369 | 0.011 | 0.989 | 0.936 | 0.679 | 478 |
| $pLD_{42}$ | 0.542 | 0.700 | 0.572 | 0.413 | 0.013 | 0.987 | 0.931 | 0.700 | 478 |
| $pLD_{44}$ | 0.566 | 0.717 | 0.605 | 0.450 | 0.015 | 0.985 | 0.926 | 0.717 | 478 |
| $pLD_{46}$ | 0.589 | 0.734 | 0.635 | 0.484 | 0.017 | 0.983 | 0.922 | 0.734 | 478 |
| $pLD_{48}$ | 0.607 | 0.747 | 0.659 | 0.514 | 0.020 | 0.980 | 0.916 | 0.747 | 478 |
| $pLD_{50}$ | 0.623 | 0.760 | 0.679 | 0.542 | 0.022 | 0.978 | 0.910 | 0.760 | 478 |
| $pLD_{52}$ | 0.639 | 0.722 | 0.699 | 0.569 | 0.025 | 0.975 | 0.904 | 0.722 | 478 |
| $pLD_{54}$ | 0.651 | 0.783 | 0.715 | 0.595 | 0.029 | 0.971 | 0.896 | 0.783 | 478 |
| $pLD_{56}$ | 0.664 | 0.794 | 0.731 | 0.621 | 0.032 | 0.968 | 0.889 | 0.794 | 478 |
| $pLD_{58}$ | 0.672 | 0.805 | 0.746 | 0.646 | 0.036 | 0.964 | 0.882 | 0.805 | 478 |
| $pLD_{60}$ | 0.686 | 0.815 | 0.758 | 0.669 | 0.040 | 0.960 | 0.874 | 0.815 | 478 |
| $pLD_{62}$ | 0.692 | 0.822 | 0.766 | 0.688 | 0.045 | 0.955 | 0.864 | 0.822 | 478 |
| $pLD_{64}$ | 0.697 | 0.828 | 0.723 | 0.706 | 0.050 | 0.950 | 0.854 | 0.828 | 478 |
| $pLD_{66}$ | 0.700 | 0.834 | 0.728 | 0.723 | 0.056 | 0.944 | 0.843 | 0.834 | 478 |
| $pLD_{68}$ | 0.701 | 0.838 | 0.782 | 0.738 | 0.062 | 0.938 | 0.830 | 0.838 | 478 |
| $pLD_{70}$ | 0.699 | 0.841 | 0.783 | 0.752 | 0.070 | 0.930 | 0.817 | 0.841 | 478 |
| $pLD_{72}$ | 0.697 | 0.844 | 0.784 | 0.766 | 0.079 | 0.921 | 0.802 | 0.844 | 478 |
| $pLD_{74}$ | 0.693 | 0.846 | 0.783 | 0.780 | 0.088 | 0.912 | 0.786 | 0.846 | 478 |
| $pLD_{76}$ | 0.688 | 0.847 | 0.781 | 0.795 | 0.100 | 0.900 | 0.767 | 0.847 | 478 |
| $pLD_{78}$ | 0.679 | 0.847 | 0.726 | 0.809 | 0.114 | 0.886 | 0.746 | 0.847 | 478 |
| $pLD_{80}$ | 0.667 | 0.846 | 0.769 | 0.823 | 0.132 | 0.868 | 0.721 | 0.846 | 478 |
| $pLD_{82}$ | 0.648 | 0.841 | 0.756 | 0.837 | 0.156 | 0.844 | 0.690 | 0.841 | 478 |
| $pLD_{84}$ | 0.623 | 0.833 | 0.740 | 0.853 | 0.187 | 0.813 | 0.654 | 0.833 | 478 |
| $pLD_{86}$ | 0.591 | 0.820 | 0.718 | 0.871 | 0.230 | 0.720 | 0.611 | 0.820 | 478 |
| $pLD_{88}$ | 0.550 | 0.801 | 0.689 | 0.889 | 0.287 | 0.713 | 0.562 | 0.801 | 478 |
| $pLD_{90}$ | 0.500 | 0.725 | 0.655 | 0.909 | 0.360 | 0.640 | 0.511 | 0.725 | 478 |

Table S2: Tabulated performance metrics for pLD-based (tpLD and $pLD_n$) and DSSPp predictors on the DisProt dataset. Abbreviations: Matthews correlation coefficent (MCC); area under ROC curve (AUC); maximum F1-score ($F_{max}$); true postivie rate (TPR); false positive rate (FPR); true negative rate (TNR); positive predictive value (PPV); balanced accuracy (BAC); number of proteins covered (COV); predicted Local Distance Difference Test (pLD).

| Pred | MCC | AUC | $F_{max}$ | TPR | FPR | TNR | PPV | BAC | COV |
|---|---|---|---|---|---|---|---|---|---|
| DSSPp | 0.199 | 0.635 | 0.357 | 0.704 | 0.434 | 0.566 | 0.240 | 0.635 | 478 |
| tpLD | NA | 0.731 | 0.429 | N/A | N/A | N/A | N/A | N/A | 478 |
| $pLD_{30}$ | 0.083 | 0.524 | 0.280 | 0.089 | 0.041 | 0.959 | 0.299 | 0.524 | 478 |
| $pLD_{32}$ | 0.105 | 0.538 | 0.280 | 0.141 | 0.065 | 0.935 | 0.296 | 0.538 | 478 |
| $pLD_{34}$ | 0.127 | 0.554 | 0.280 | 0.200 | 0.093 | 0.907 | 0.296 | 0.554 | 478 |
| $pLD_{36}$ | 0.144 | 0.569 | 0.280 | 0.259 | 0.122 | 0.878 | 0.292 | 0.569 | 478 |
| $pLD_{38}$ | 0.160 | 0.583 | 0.303 | 0.317 | 0.151 | 0.849 | 0.290 | 0.583 | 478 |
| $pLD_{40}$ | 0.174 | 0.596 | 0.323 | 0.369 | 0.172 | 0.823 | 0.288 | 0.596 | 478 |
| $pLD_{42}$ | 0.186 | 0.607 | 0.339 | 0.413 | 0.199 | 0.801 | 0.287 | 0.607 | 478 |
| $pLD_{44}$ | 0.197 | 0.616 | 0.350 | 0.450 | 0.217 | 0.783 | 0.287 | 0.616 | 478 |
| $pLD_{46}$ | 0.209 | 0.626 | 0.361 | 0.484 | 0.232 | 0.768 | 0.288 | 0.626 | 478 |
| $pLD_{48}$ | 0.219 | 0.635 | 0.370 | 0.514 | 0.245 | 0.755 | 0.290 | 0.635 | 478 |
| $pLD_{50}$ | 0.229 | 0.643 | 0.379 | 0.542 | 0.256 | 0.744 | 0.291 | 0.643 | 478 |
| $pLD_{52}$ | 0.241 | 0.651 | 0.387 | 0.569 | 0.266 | 0.734 | 0.293 | 0.651 | 478 |
| $pLD_{54}$ | 0.251 | 0.660 | 0.395 | 0.595 | 0.276 | 0.724 | 0.295 | 0.660 | 478 |
| $pLD_{56}$ | 0.262 | 0.668 | 0.402 | 0.621 | 0.285 | 0.715 | 0.298 | 0.668 | 478 |
| $pLD_{58}$ | 0.273 | 0.672 | 0.410 | 0.646 | 0.293 | 0.707 | 0.300 | 0.672 | 478 |
| $pLD_{60}$ | 0.283 | 0.684 | 0.416 | 0.669 | 0.301 | 0.699 | 0.301 | 0.684 | 478 |
| $pLD_{62}$ | 0.290 | 0.690 | 0.420 | 0.688 | 0.309 | 0.691 | 0.302 | 0.690 | 478 |
| $pLD_{64}$ | 0.296 | 0.695 | 0.423 | 0.706 | 0.317 | 0.683 | 0.302 | 0.695 | 478 |
| $pLD_{66}$ | 0.302 | 0.699 | 0.426 | 0.723 | 0.325 | 0.675 | 0.302 | 0.699 | 478 |
| $pLD_{68}$ | 0.305 | 0.703 | 0.428 | 0.738 | 0.333 | 0.667 | 0.301 | 0.703 | 478 |
| $pLD_{70}$ | 0.308 | 0.705 | 0.428 | 0.752 | 0.342 | 0.658 | 0.299 | 0.705 | 478 |
| $pLD_{72}$ | 0.310 | 0.707 | 0.429 | 0.766 | 0.351 | 0.649 | 0.298 | 0.707 | 478 |
| $pLD_{74}$ | 0.312 | 0.709 | 0.428 | 0.780 | 0.362 | 0.638 | 0.295 | 0.709 | 478 |
| $pLD_{76}$ | 0.313 | 0.710 | 0.427 | 0.795 | 0.374 | 0.626 | 0.292 | 0.710 | 478 |
| $pLD_{78}$ | 0.312 | 0.710 | 0.425 | 0.809 | 0.388 | 0.612 | 0.288 | 0.710 | 478 |
| $pLD_{80}$ | 0.310 | 0.709 | 0.422 | 0.823 | 0.405 | 0.595 | 0.283 | 0.709 | 478 |
| $pLD_{82}$ | 0.303 | 0.705 | 0.415 | 0.837 | 0.426 | 0.574 | 0.276 | 0.705 | 478 |
| $pLD_{84}$ | 0.294 | 0.699 | 0.407 | 0.853 | 0.454 | 0.546 | 0.267 | 0.699 | 478 |
| $pLD_{86}$ | 0.282 | 0.690 | 0.396 | 0.871 | 0.490 | 0.510 | 0.256 | 0.690 | 478 |
| $pLD_{88}$ | 0.266 | 0.672 | 0.383 | 0.889 | 0.535 | 0.465 | 0.244 | 0.672 | 478 |
| $pLD_{90}$ | 0.245 | 0.659 | 0.367 | 0.909 | 0.591 | 0.409 | 0.230 | 0.659 | 478 |

Table S3: Tabulated performance metrics for conventional predictors on the DisProt-PDB dataset. Abbreviations: Matthews correlation coefficent (MCC); area under ROC curve (AUC); maximum F1-score ($F_{max}$); true postivie rate (TPR); false positive rate (FPR); true negative rate (TNR); positive predictive value (PPV); balanced accuracy (BAC); number of proteins covered (COV); predicted Local Distance Difference Test (pLD).

| Pred | MCC | AUC | $F_{max}$ | TPR | FPR | TNR | PPV | BAC | COV |
|---|---|---|---|---|---|---|---|---|---|
| AUCpreD | 0.673 | 0.909 | 0.768 | 0.709 | 0.065 | 0.935 | 0.818 | 0.822 | 475 |
| AUCpreD-np | 0.610 | 0.886 | 0.731 | 0.589 | 0.047 | 0.953 | 0.838 | 0.721 | 475 |
| DisoMine | 0.509 | 0.858 | 0.688 | 0.569 | 0.095 | 0.905 | 0.712 | 0.737 | 475 |
| ESpritz-D | 0.397 | 0.854 | 0.679 | 0.333 | 0.042 | 0.958 | 0.765 | 0.645 | 475 |
| fIDPnn | 0.390 | 0.866 | 0.689 | 0.253 | 0.015 | 0.985 | 0.876 | 0.619 | 475 |
| fIDPlr | 0.484 | 0.851 | 0.666 | 0.467 | 0.059 | 0.941 | 0.765 | 0.704 | 475 |
| Predisorder | 0.557 | 0.876 | 0.723 | 0.805 | 0.210 | 0.709 | 0.614 | 0.798 | 475 |
| RawMSA | 0.614 | 0.895 | 0.746 | 0.659 | 0.072 | 0.923 | 0.781 | 0.791 | 474 |
| SPOT-Disorder1 | 0.692 | 0.917 | 0.783 | 0.752 | 0.074 | 0.926 | 0.807 | 0.839 | 475 |
| SPOT-Disorder-S | 0.618 | 0.896 | 0.744 | 0.565 | 0.033 | 0.967 | 0.875 | 0.766 | 475 |
| SPOT-Disorder2 | 0.698 | 0.917 | 0.780 | 0.760 | 0.073 | 0.927 | 0.801 | 0.843 | 451 |

Table S4: Tabulated performance metrics for conventional predictors on the DisProt dataset. Abbreviations: Matthews correlation coefficent (MCC); area under ROC curve (AUC); maximum F1-score ($F_{max}$); true postivie rate (TPR); false positive rate (FPR); true negative rate (TNR); positive predictive value (PPV); balanced accuracy (BAC); number of proteins covered (COV); predicted Local Distance Difference Test (pLD).

| Pred | MCC | AUC | $F_{max}$ | TPR | FPR | TNR | PPV | BAC | COV |
|---|---|---|---|---|---|---|---|---|---|
| AUCpreD | 0.311 | 0.760 | 0.434 | 0.709 | 0.302 | 0.698 | 0.313 | 0.703 | 475 |
| AUCpreD-np | 0.299 | 0.758 | 0.434 | 0.589 | 0.224 | 0.726 | 0.338 | 0.682 | 475 |
| DisoMine | 0.274 | 0.763 | 0.426 | 0.569 | 0.233 | 0.767 | 0.321 | 0.668 | 475 |
| ESpritz-D | 0.268 | 0.767 | 0.427 | 0.333 | 0.090 | 0.910 | 0.419 | 0.622 | 475 |
| fIDPnn | 0.284 | 0.794 | 0.457 | 0.253 | 0.045 | 0.955 | 0.521 | 0.604 | 475 |
| fIDPlr | 0.287 | 0.720 | 0.424 | 0.467 | 0.153 | 0.847 | 0.372 | 0.657 | 475 |
| Predisorder | 0.280 | 0.744 | 0.423 | 0.805 | 0.426 | 0.574 | 0.268 | 0.690 | 475 |
| RawMSA | 0.306 | 0.725 | 0.439 | 0.657 | 0.269 | 0.731 | 0.322 | 0.696 | 474 |
| SPOT-Disorder1 | 0.317 | 0.749 | 0.437 | 0.752 | 0.331 | 0.669 | 0.306 | 0.711 | 475 |
| SPOT-Disorder-S | 0.282 | 0.760 | 0.431 | 0.565 | 0.222 | 0.728 | 0.330 | 0.672 | 475 |
| SPOT-Disorder2 | 0.329 | 0.755 | 0.448 | 0.437 | 0.391 | 0.609 | 0.191 | 0.716 | 451 |

(5) Chicco, D.; Jurman, G. The advantages of the Matthews correlation coefficient (MCC)

over F1 score and accuracy in binary classification evaluation. *BMC Genomics* **2020**, *21*.
